# Spontaneous modulations of high frequency cortical activity

**DOI:** 10.1101/2021.04.15.440025

**Authors:** Hiroya Ono, Masaki Sonoda, Brian H. Silverstein, Kaori Sonoda, Takafumi Kubota, Aimee F. Luat, Robert Rothermel, Sandeep Sood, Eishi Asano

**Author notes:** Corresponding Author: Eishi Asano, M.D., Ph.D., M.S. (C.R.D.S.A.) Address: Division of Pediatric Neurology, Children’s Hospital of Michigan, Wayne State University. 3901 Beaubien St., Detroit, MI, 48201, USA. Equal contribution (Hiroya Ono, Masaki Sonoda).

## Abstract

**Objective:** We clarified the clinical and mechanistic significance of physiological modulations of high-frequency broadband cortical activity associated with spontaneous saccadic eye movements during a resting state.

**Methods:** We studied 30 patients who underwent epilepsy surgery following extraoperative electrocorticography and electrooculography recordings. We determined whether high-gamma activity at 70-110 Hz *<u>preceding saccade onset</u>* would predict upcoming ocular behaviors. We assessed how accurately the model incorporating saccade-related high-gamma modulations would localize the primary visual cortex defined by electrical stimulation.

**Results:** The whole-brain level dynamic atlas demonstrated transient high-gamma suppression in the striatal region before saccade onset and high-gamma augmentation subsequently involving the widespread posterior brain regions. More intense striatal high-gamma suppression predicted the upcoming saccade directed to the ipsilateral side and lasting longer in duration. The bagged-tree-ensemble model demonstrated that intense saccade-related high-gamma modulations localized the visual cortex with an accuracy of 95%.

**Conclusions:** We successfully animated the neural dynamics supporting saccadic suppression, a principal mechanism minimizing the perception of blurred vision during rapid eye movements. The primary visual cortex *per se* may prepare *<u>actively in advance</u>* for massive image motion expected during upcoming prolonged saccades.

**Significance:** Measuring saccade-related electrocorticographic signals may help localize the visual cortex and avoid misperceiving physiological high-frequency activity as epileptogenic.

**Highlights:**

- The whole-brain level dynamic atlas animated spontaneous high gamma modulations associated with saccadic eye movements.
- Preceding high gamma activity in the striatal cortex predicted the direction and duration of the upcoming saccades.
- Saccade-related high-gamma modulations localized the stimulation-defined visual cortex with an accuracy of 95%.

## 1. Introduction

Our stable visual perception is, in part, secured by a transient reduction of visual sensitivity during a rapid eye movement, and this brain function is often referred to as ‘saccadic suppression’ (Duffy and Lombroso, 1968; Matin, 1974; Thiele *et al*., 2002). Saccadic suppression effectively keeps us unaware of image motion across the retina during fast eye movements taking place spontaneously and inevitably every second during wakefulness (Schiller and Tehovnik, 2001). The association between impaired saccadic suppression and amblyopia has been previously suggested (Martinez-Conde *et al*., 2013). Single-neuron studies of non-human primates reported that saccade-related neuronal suppression, reflected by a reduced firing rate during saccades, involved the superior colliculus in the midbrain as well as the striatal and extra-striatal visual pathways, including the middle-temporal visual area (Duffy and Burchfiel, 1975; Zanos *et al*. 2016; Berman *et al*., 2017). Previous intracranial EEG (iEEG) studies of 10-12 patients with focal epilepsy successfully demonstrated that spontaneous saccades resulted in transient suppression of neuronal activity at lateral and ventral occipital areas (Uematsu *et al*., 2013; Golan *et al*., 2017). The large-scale spatial gradient of suppressive function was not fully delineated, mainly because the spatial extent of invasive signal sampling is inevitably limited in an individual patient (Uematsu *et al*., 2013; Golan *et al*., 2017). The present study has overcome this spatial limitation by combining iEEG measures at 2,290 nonepileptic electrode sites from 30 patients and generated a four-dimensional (4D) functional atlas animating the neuronal dynamics around spontaneous saccades in the three-dimensional (3D) space. Such a rare and unique iEEG dataset allowed us to comprehensively model the spatiotemporal dynamics of neuronal modulations in the perisaccadic period.

We designed this study to visualize what cortical structure would actively initiate saccade-related neuronal suppression and how neuronal suppression would propagate. Based on the observations in previous studies of non-human primates (Duffy and Burchfiel, 1975; Zanos *et al*. 2016; Berman *et al*., 2017), primates, we expect that saccade-related neuronal suppression at the cortical level would take place initially in the striatal region and propagate to the lateral occipital and fusiform regions. **[Aim 1]** We tested the specific hypothesis that, independently of the effects of patient demographics and epilepsy-related variables, the saccade-related neuronal suppression would be most intense and lingering in the posterior portion within the striatal cortex. This hypothesis is based on the notion that the human striatal cortex has an *anterior-to-posterior* topographic gradient in support of *peripheral-to-central* receptive fields (Horton and Hoyt, 1991; Brewer *et al*., 2005; Ruff *et al*., 2006; Yoshor *et al*., 2007). Behavioral studies of healthy humans previously suggested that saccade-related reduction of visual sensitivity involves the central/near-foveal more than the peripheral field (Matin, 1974; Bridgeman and Fisher, 1990).

**[Aim 2]** We tested the hypothesis that the direction of given saccades would alter the magnitude of saccade-related neuronal suppression. It is plausible to expect that our object recognition in a natural scene would be optimized by reducing visual sensitivity in an unattended hemifield (Wurtz, 2008; Cavanaugh *et al*., 2016). A human behavior study reported that saccadic suppression in a hemifield was modestly dependent on the direction of eye movement (Osaka, 1987). A single-neuron study of non-human primates reported that saccade-related neuronal suppression was grossly agnostic to the direction of eye movement and took place in the primary visual cortex supporting both hemifields contralateral and ipsilateral to a given eye movement (Berman *et al*., 2017).

**[Aim 3]** The present study further tested the hypothesis that intense neuronal suppression preceding saccade onset would predict a prolonged duration of the upcoming saccade. We believe that this analysis would provide evidence that the cerebral cortex *per se* actively prepares it in advance for massive image motion expected during long saccades; such anticipation-related preparation is referred to as ‘efferent copy’ or ‘corollary discharge’ (Bridgeman *et al*., 1994). Single-neuron studies of non-human primates, as well as behavioral and functional MRI studies of healthy humans, suggest that either or both striatal and extra-striatal visual pathways may be responsible for the efferent copy (Kleiser *et al*., 2004; Vallines and Greenlee, 2006; Chahine and Krekelberg, 2009; Martinez-Conde *et al*., 2013). Previous iEEG studies of patients with epilepsy likewise reported that striatal or extra-striatal visual areas were transiently suppressed *<u>during</u>* saccades (Uematsu *et al*., 2013; Golan *et al*., 2017). However, these iEEG studies were not designed to clarify whether the magnitude of *<u>preceding neuronal suppression</u>* would predict the subsequent saccade behaviors of given individuals on a trial-by-trial basis.

The present study quantified the amplitude of broadband activity at high-gamma range (70-110 Hz) time-locked to saccade onset and offset to assess the association between perisaccadic neural modulations and saccade behaviors. High-gamma amplitude is an excellent summary measure of event-related neuronal modulations (Crone *et al*., 2011; Nakai *et al*., 2017). An increase in high-gamma amplitude is highly associated with an increase in firing rate (Ray *et al*., 2008), hemodynamic response (Scheeringa *et al*., 2011), and glucose metabolism (Nishida *et al*., 2008). Conversely, reduced high-gamma amplitude is associated with a reduction in firing rate and hemodynamic response (Shmuel *et al*., 2006). Because a single high-gamma cycle is as short as 14 ms, one can assess the neural dynamics with a temporal resolution of tens of milliseconds’ order. The clinical significance of local event-related high-gamma modulations is better understood than that of low-frequency band activities (e.g., beta). Resection of sites showing event-related high-gamma augmentation frequently results in a functional deficit requiring rehabilitation therapy (Kojima *et al*., 2013). Electrical stimulation of task-related high-gamma sites often interferes with executing a given task (Arya *et al*., 2018*a*).

**[Aim 4]** The present study finally tested the hypothesis that task-free high-gamma modulations during saccades would accurately localize the primary visual cortex defined by the gold-standard electrical stimulation mapping (ESM). Investigators previously reported that the occurrence rate and amplitude of interictal high-frequency broadband activities were increased at healthy occipital areas and speculated that such neural modulations might be associated with lower-order sensory processing (Nagasawa et al., 2012; Alkawadri et al., 2014; Frauscher et al., 2018; Motoi et al., 2019). We hope our multimodality analysis has provided definitive evidence that measurement of spontaneous saccade-related high-gamma modulations helps localize the primary visual areas. The results are expected to bring a meaningful insight into the characterization of task-free high-frequency cortical activities in epilepsy presurgical evaluation.

## 2. Methods

### 2.1. Patients

We studied a total of 30 patients with drug-resistant focal epilepsy (age range: 5-20 years; 16 females; **Table 1**) who satisfied the following criteria. The inclusion criteria included [a] simultaneous video-iEEG and electrooculography (EOG) recording as part of our routine presurgical evaluation at Children’s Hospital of Michigan in Detroit from Dec 2008 to July 2018, [b] iEEG sampling from the occipital lobe, and [c] at least 80 events of spontaneous saccades to both left and right directions available during EOG recording. The exclusion criteria included [a] age of less than four [b] presence of seizure onset zone (SOZ), interictal spike discharges, or structural lesions involving the occipital lobe (Asano *et al*., 2009; Motoi *et al*., 2019), [c] visual field deficits prior to surgery, and [d] history of previous epilepsy surgery. This study was approved by the Wayne State University Institutional Review Board, and informed consent in writing was obtained from the patients or guardians of patients.

**Table 1:** Patient profile.

|  |  |
| --- | --- |
| Patient profile. |  |
| Number of patients | 30 |
| Mean of age in years (range) | 12.0 (5 - 20) |
| Proportion of male (%) | 46.7 |
| Sampled hemisphere (%) |  |
| Left | 50.0 |
| Right | 50.0 |
| Seizure onset zone (%) |  |
| involving the frontal lobe | 23.3 |
| involving the temporal lobe | 43.3 |
| involving the parietal lobe | 23.3 |
| Proportion of patients with an MRI lesion (%) | 76.7 |
| Mean number of antiepileptic drugs (SD) | 1.9 (0.61) |

SD = Standard deviation.

### 2.2. iEEG data acquisition

We previously reported the iEEG data acquisition protocol (Nakai *et al*., 2017). We surgically implanted platinum grid/strip electrodes (3 mm exposed diameter and 10 mm center-to-center distance) into the affected hemisphere’s subdural space. We clinically determined the spatial extent of subdural electrode placement to localize the boundary between the epileptogenic zone to be removed and the functionally-important areas to be preserved (Asano *et al*., 2009). We acquired video-iEEG recordings continuously at the bedside using a 192-channel Nihon Kohden Neurofax 1100A Digital System (Nihon Kohden America Inc, Foothill Ranch, CA, USA). We set the iEEG sampling frequency at 1,000 Hz and the amplifier band-pass at 0.016 to 300 Hz. We set the averaged voltage of iEEG signals derived from the fifth and sixth intracranial electrodes of the amplifier as the original reference and re-montaged the iEEG signals to a common average reference (Lesser *et al*., 2010; Nariai *et al*., 2011). Thereby, we excluded channels classified as SOZ, showing interictal spikes, affected by structural lesions, or artifacts from the common average reference and further analyses described below. Thus, a total of 2,290 artifact-free nonepileptic channels (77.1 electrode sites per patient) were available for the iEEG analysis (**Fig. 1A**).

**Fig. 1.**
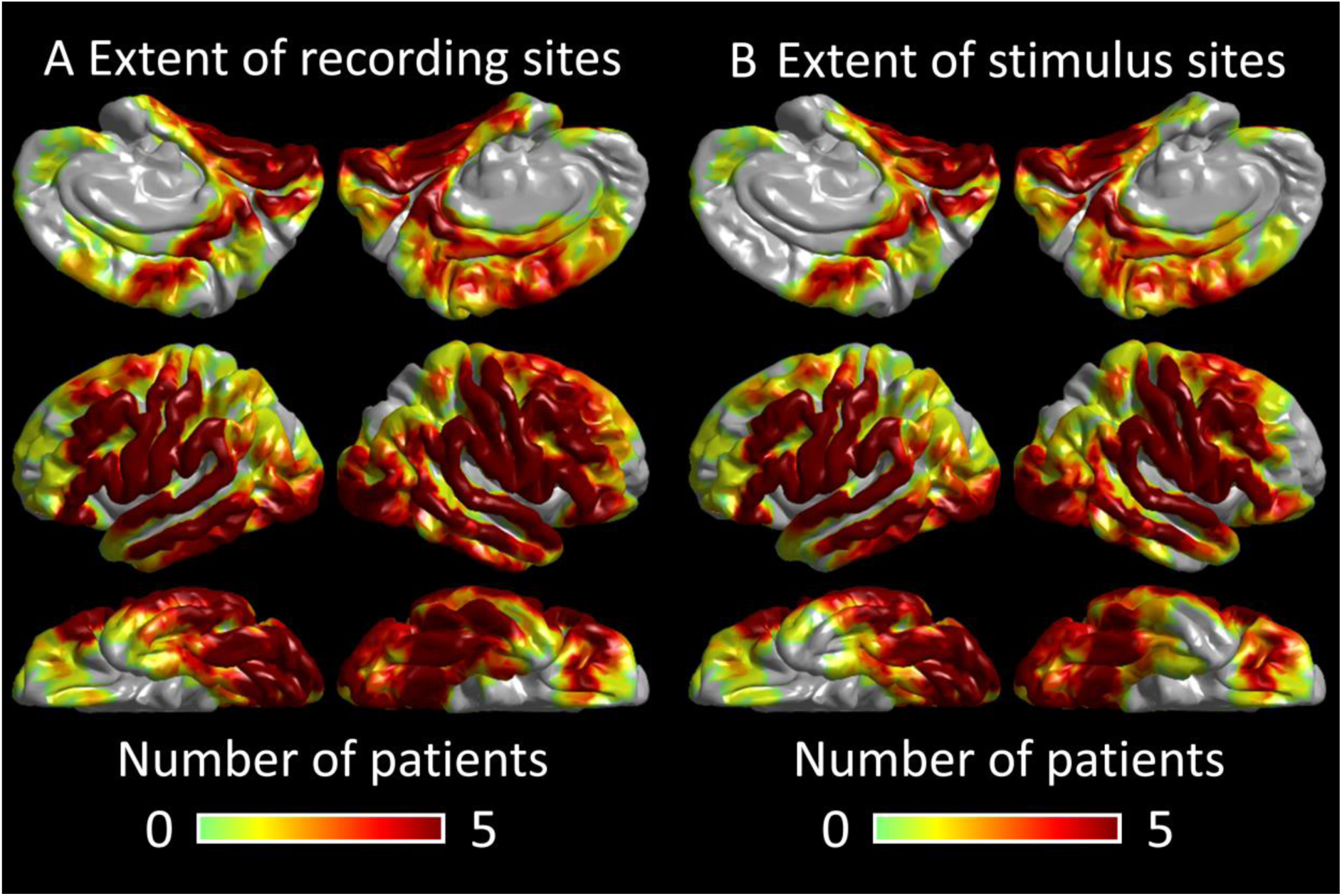
The extent of non-epileptic electrode sites. (**A**) The spatial distribution of the analyzed electrodes on each hemisphere is presented on a color-mapped 3D brain surface image. The color of each site indicates the number of patients available at a given spatial point. Of the total 2,290 non-epileptic artifact-free electrode sites, the number of electrode sites within each region of interest (ROI) is as follows. Striatal: 127 electrode sites. Lateral-occipital: 140. Fusiform: 104. Frontal-eye field (FEF): 77. (**B**) The number of electrode sites which were stimulated for cortical mapping is presented. Of the total 1,903 electrode sites, the number of electrode sites within each ROI is as follows. Striatal: 127. Lateral-occipital: 140. Fusiform: 104.

### 2.3. 3D MRI data processing

We generated a 3D surface image with implanted electrodes displayed directly on the cortical surface using a preoperative T1-weighted spoiled gradient-recalled echo sequence MRI and a CT image immediately following the placement of intracranial electrodes (Nakai *et al*., 2017; Stolk *et al*., 2018). Using intraoperative photographs, we confirmed the spatial accuracy of the coregistration of MR surface image and subdural electrodes (Pieters *et al*., 2013). We subsequently normalized electrode sites of a given patient to the FreeSurfer coordinates (http://surfer.nmr.mgh.harvard.edu) to enable the group-level visualization and analysis. We then employed an automatic parcellation of cortical gyri at both individual and normalized surfaces to assign each electrode site a region of interest (ROI) (Desikan *et al*., 2006; Nakai *et al*., 2019; **Fig. 2; Table S1**). The Desikan FreeSurfer atlas did not explicitly define the spatial extent of the frontal eye field (FEF). As previously performed (Sugiura *et al*., 2020), the present study defined the FEF as the regions at which 50-Hz stimulation consistently resulted in a forced eye deviation in ≥5% of the 84 adult patients included in our previous ESM study (**Fig. 2**).

**Fig. 2.**
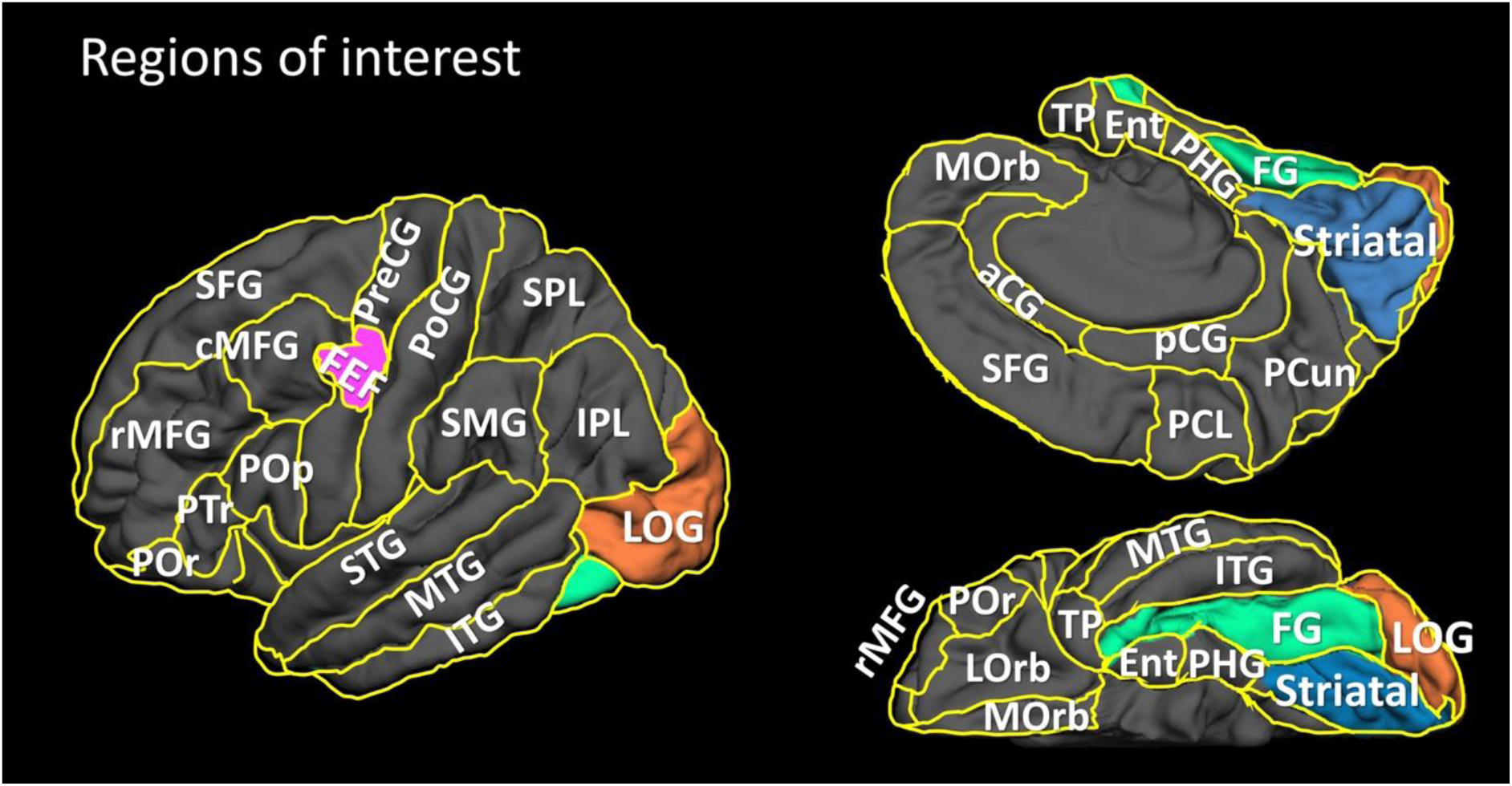
Regions of interest (ROIs). All ROIs in the present study are presented (Nakai et al., 2019; Sugiura et al., 2020). Color-coded ROIs include striatal region (light blue), lateral-occipital gyrus (LOG; orange), fusiform gyrus (FG; green), and frontal-eye field (FEF; purple). The other ROIs are anterior cingulate gyrus (aCG), caudal middle frontal gyrus (cMFG), entorhinal gyrus (Ent), inferior parietal lobule (IPL), inferior-temporal gyrus (ITG), lateral orbitofrontal gyrus (LOrb), medial orbitofrontal gyrus (MOrb), middle-temporal gyrus (MTG), paracentral gyrus (PCL), parahippocampal gyrus (PHG), postcentral gyrus (PoCG), pars opercularis of the inferior-frontal gyrus (Pop), pars orbitalis of the inferior-frontal gyrus (POr), pars triangularis of the inferior-frontal gyrus (PTr), posterior cingulate gyrus (pCG), precentral gyrus (PreCG), precuneus (PCun), rostral middle-frontal gyrus (rMFG), superior frontal gyrus (SFG), supramarginal gyrus (SMG), superior parietal lobule (SPL), superior-temporal gyrus (STG), and temporal pole (TP).

### 2.4. Marking of saccade events

As part of our routine presurgical evaluation, we placed EOG electrodes 2.5 cm below and 2.5 cm lateral to the left and right outer canthi and monitored spontaneous eye movements during the video-iEEG recording (Uematsu *et al*., 2013; Kambara *et al*., 2018). With video assistance, we visually identified and marked the onset and offset of at least 80 events of spontaneous horizontal saccades in each direction (**Fig. 3**) that took place during a task-free wakeful resting state. We excluded periods during cognitive tasks (Nakai *et al*., 2019) or within two hours following seizures from analyses. We defined a saccade as an abrupt and sharp ocular movement reflected by an EOG amplitude change of ≥40 μV with a duration ranging from 25 to 700 ms on iEEG with a time constant set at 0.1 s (Baloh *et al*., 1975; Kaufman and Abel, 1986; Martinez-Conde *et al*., 2009; Uematsu *et al*., 2013). A longer saccade duration is suggested to reflect a larger eye movement in general (Bahill *et al*., 1981; Findlay and Gilchrist, 2003; Qing and Kapoula, 2004). In the present study of spontaneous gross saccades during task-free, naturalistic, and exploratory viewing, we did not mark a saccadic event with its duration ranging below 25 ms. Reliable identification of such miniature saccades (also known as micro/fixational saccades) would require an eye-tracker with a high sampling rate (Yuval-Greenberg *et al*., 2008; Dimigen *et al*., 2009). The maximum duration of micro-saccades was reported to be 300 ms in a study of non-human primates instructed to detect visual stimuli moving on a computer screen (Herrington *et al*., 2009). In contrast, our patients were in a natural patient room environment and might have had large eye movements. Anticipatory gross saccades were previously reported to last up to 1,500 ms in duration (Kaufman and Abel, 1986).

**Fig. 3.**
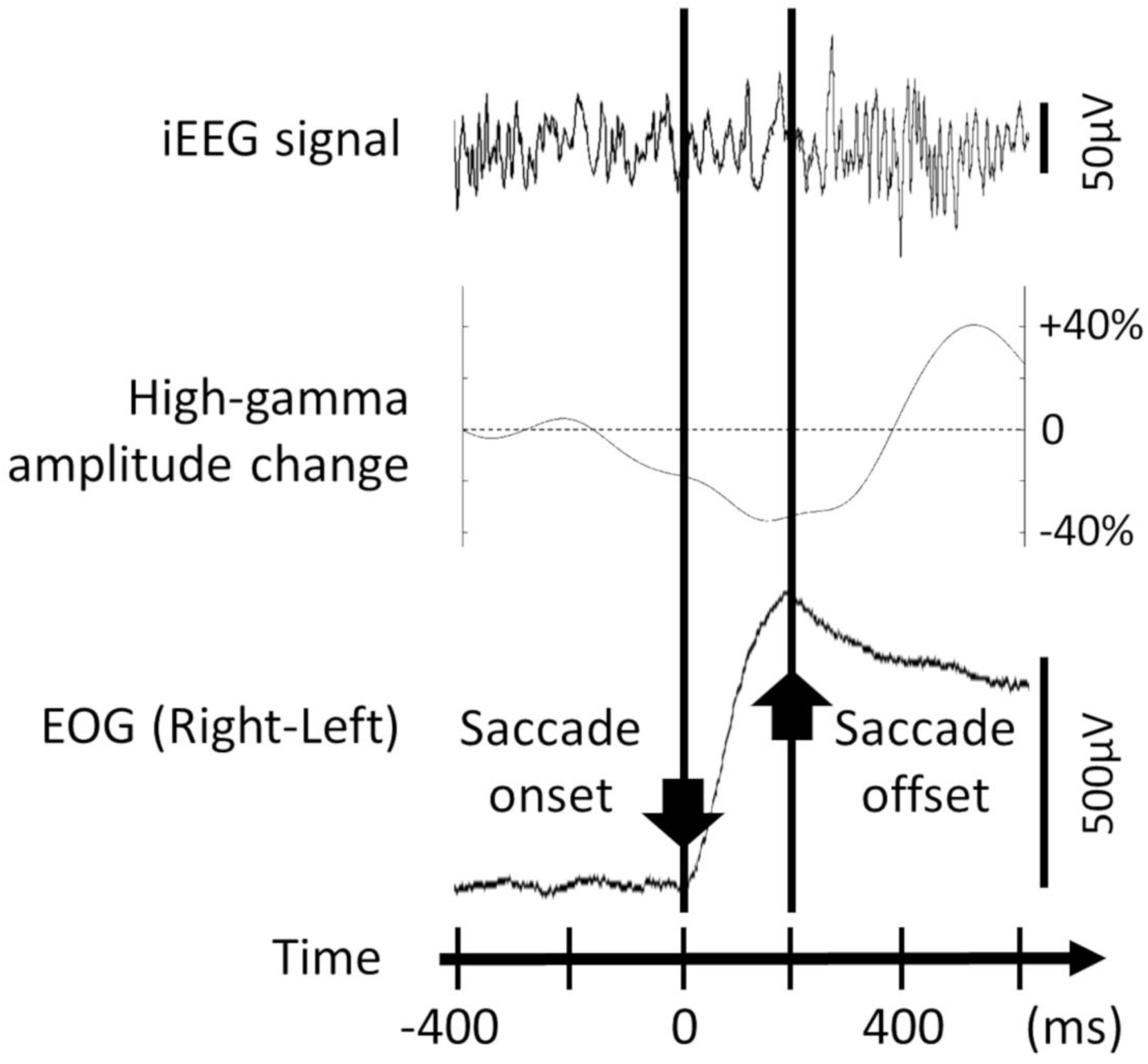
Marking of saccade onset and offset in a 14-year-old girl with left temporal lobe epilepsy. Upper trace: Intracranial EEG (iEEG) trace at a striatal site (High-cut filter: 300 Hz. Time constant: 0.003 s). Middle trace: Dynamics of striatal high-gamma amplitude modulations compared to that during the baseline period between 600 and 200 ms prior to the saccade onset. Lower trace: Electrooculography (EOG) trace determined the timing of the saccade onset and offset (High-cut filter: 300 Hz. Time constant: 10 s).

### 2.5. Time-frequency analysis to determine the dynamics of peri-saccadic high-gamma modulations

Time-frequency analysis determined when and where high-gamma band amplitude was augmented and attenuated compared to that during the baseline/reference period (specified in detail below). The complex demodulation method transformed iEEG signals into 5-ms/10-Hz time-frequency bins during a 3,000-ms time window (between 1,000 ms before and 2,000 ms after each saccade onset and offset; Papp and Ktonas, 1977; Hoechstetter *et al*., 2004). This method, employing a low-pass finite impulse response (FIR) filter of Gaussian shape, was effectively equivalent to a Gabor transform. The resolution of the time-frequency analysis was ±7.9 ms and ±14.2 Hz (Hoechstetter *et al*., 2004). At given 5 ms epochs at each electrode site, we computed the percent change of high-gamma amplitude in the 70-110 Hz range relative to the mean amplitude during the baseline period.

One cannot assume a baseline period completely free from another saccade event because exploratory saccadic eye movements inevitably, spontaneously and unpredictably occur every second. Thus, we determined peri-saccadic high-gamma modulations using the following three different baselines. [**Fixed baseline approach**] The baseline was defined as the average across high-gamma amplitudes during the 400-ms period at 200-600 ms prior to saccade onset (**Fig. S1**; Nakai *et al*., 2019). [**Jittered baseline approach**] The baseline was defined as the average across high-gamma amplitudes during a randomly-selected 200-ms period between saccade onset and 800 ms before saccade onset (**Fig. S1**). [**Normalization approach**] The baseline period was defined as the entire 3,000-ms period between 1,000 ms before and 2,000 ms after saccade onset. At a given 5-ms time window and each site, we computed ‘high-gamma z-score’, reflecting how much high-gamma amplitude deviated from the mean during the baseline period (**Fig. S1**). The Spearman’s rank test determined whether the spatial distribution of high-gamma modulations during the post-saccade 400-ms period (as rated by ranking; Mitsuhashi *et al*., 2020) would be similar between the fixed and jittered baseline approaches as well as between the fixed baseline and normalization approaches. If the spatial distribution of saccade-related high-gamma activity was similar across these approaches, we reported the high-gamma measures based on the fixed baseline approach alone.

### 2.6. Visualization of peri-saccadic high-gamma modulations

We initially presented the dynamics of high-gamma activity time-locked to saccade onset and offset at given ROIs (**Fig. 2**). The one-sample t-test, followed by the false discovery rate (FDR) correction, determined the time window in which the averaged high-gamma amplitude at a given ROI differed from that during the baseline period (**Fig. 4**). The corrected significance level α was set to 0.05. We subsequently presented the dynamics of high-gamma modulations, at the group level, on the averaged FreeSurfer pial surface image with each electrode site spatially normalized to template space, as previously reported (Nakai *et al*., 2017; Sugiura *et al*., 2020; **Video S1; Fig. 5**).

**Fig. 4.**
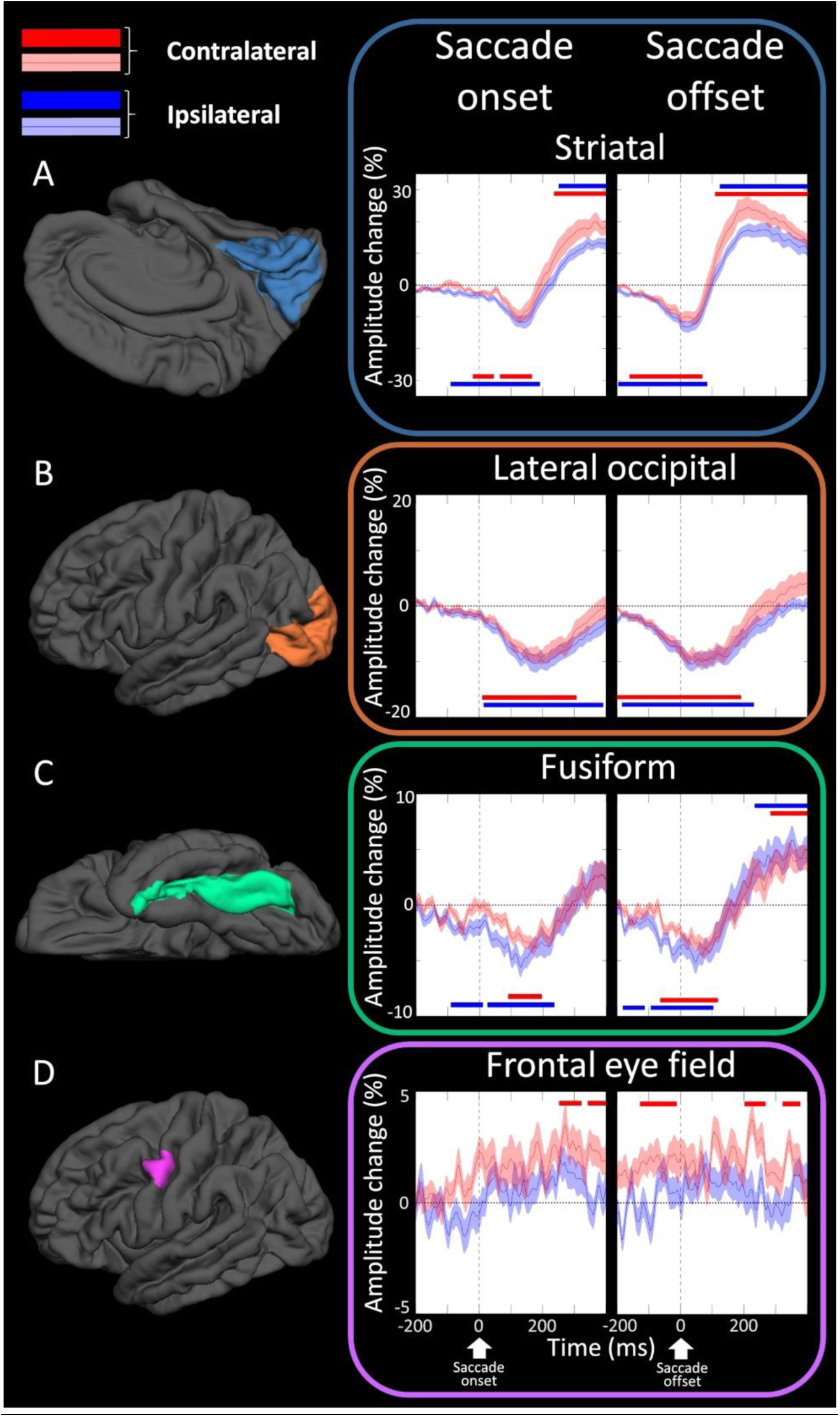
Saccade-related high-gamma modulations. (**A**) Striatal region (light blue). (**B**) Lateral-occipital region (orange). (**C**) Fusiform region (green). (**D**) Frontal-eye field (FEF; purple). The plots demonstrate the temporal dynamics of high-gamma amplitude time-locked to saccade onset and offset. Red line: High-gamma dynamics during saccades directed to the side contralateral to the sampled hemisphere (standard error shades are provided). Blue line: High-gamma dynamics during ipsilateral saccades. Horizontal bars denote the timing when high-gamma augmentation (upper bar) or attenuation (lower bar) reached the significance at least for 40 ms.

**Fig. 5.**
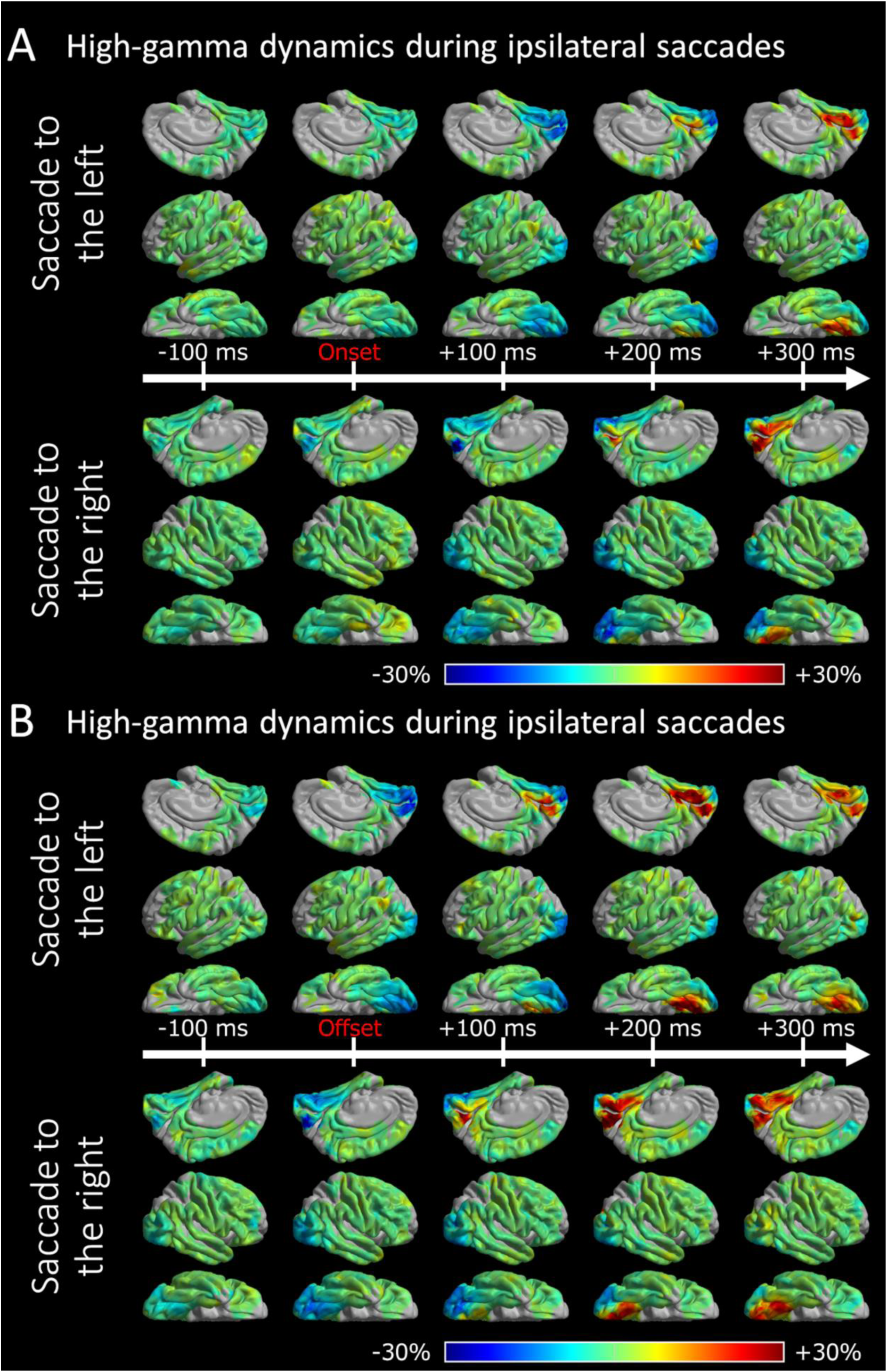
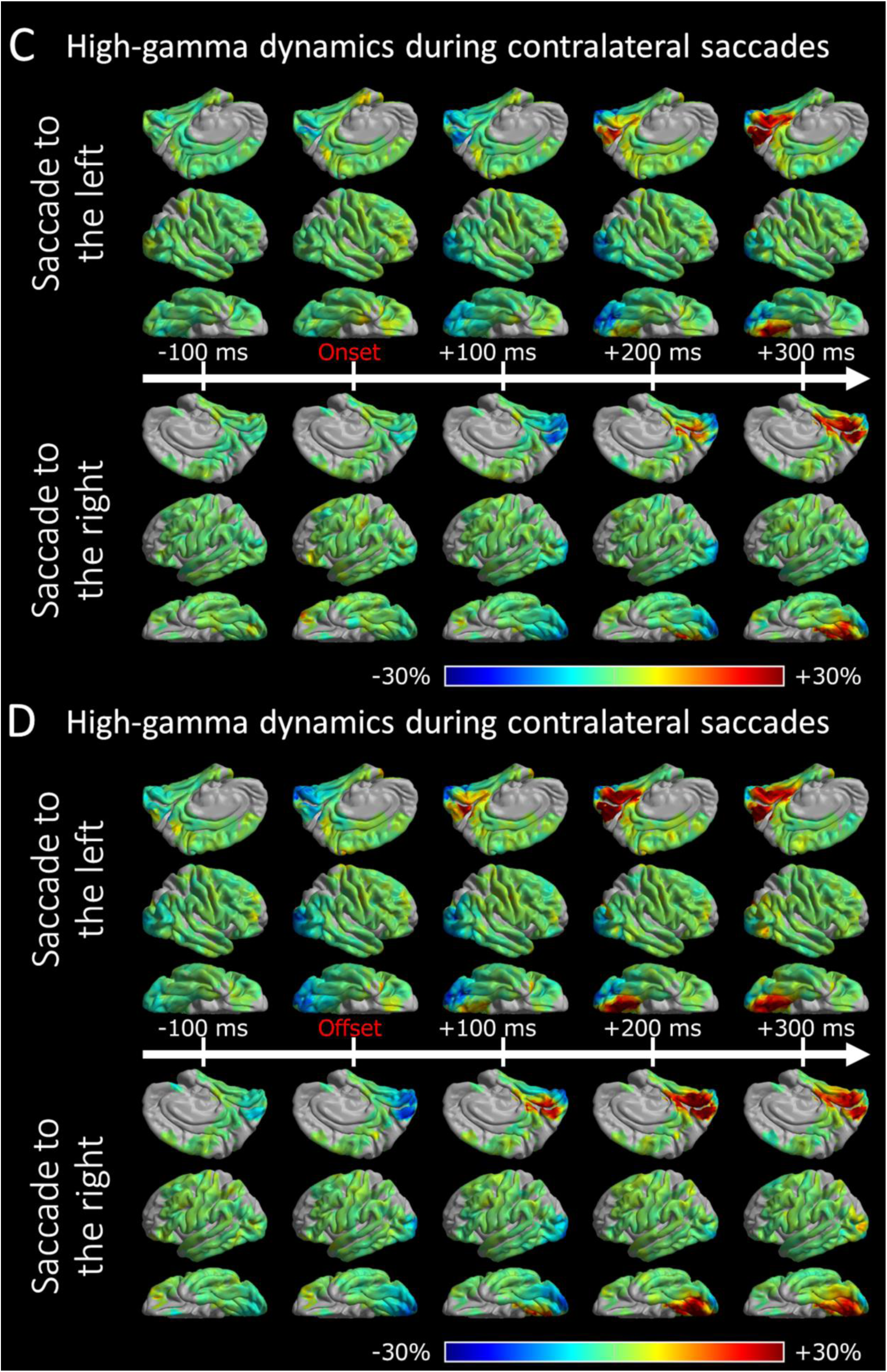
The spatiotemporal dynamics of peri-saccadic high-gamma modulations. The video snapshots demonstrate the group-level percent change of saccade-related high-gamma activity relative to the baseline period (200-600 ms prior to saccade onset) derived from all 30 patients. (**A-B**) Saccades directed to the side ipsilateral to the iEEG sampling hemisphere. (**A**) 0 ms: saccade onset. (**B**) 0 ms: saccade offset. (**C-D**) Contralateral saccades. (**C**) 0 ms: saccade onset. **(D**) 0 ms: saccade offset. High-gamma activity is displayed at sites where at least two patients’ data were available.

For interested readers, we presented the dynamics of peri-saccadic modulations of alpha (8-12 Hz) and beta (16-28 Hz) activities (**Videos S2-S3**). Based on the previous iEEG studies (Crone *et al*., 2011; Uematsu *et al*., 2013), we expected alpha and beta activities to be attenuated with high-gamma activity simultaneously augmented. Conversely, alpha and beta activities were expected to be augmented when high-gamma activity was attenuated. Compared to high-gamma activity, alpha or beta activity might be less suitable to evaluate the rapid neuronal dynamics, partly because the period of a single cycle of 10-Hz oscillations is 100 ms.

### 2.7. Electrical stimulation mapping (ESM) to localize the primary visual cortex

We performed ESM as part of clinical management with the method identical to that reported previously (Nakai *et al*., 2017, 2018). The spatial extent of ESM was purely based on the clinical need, and we found that 83.1% of the 2,290 electrode sites (63.4 electrode sites per patient) were tested in the present study (**Fig. 1B)**. A given patient was aware of the timing of stimulation but not the location. We terminated stimulation as soon as a given patient reported a percept or exhibited a symptom. The neuropsychologist (R.R.), not being aware of the results of event-related high-gamma activity, asked a given patient to characterize a percept induced by stimulation. We initially set the stimulus intensity at 3 mA and frequency at 50 Hz. We then increased the intensity until a symptom or after-discharge was observed. We did not increase the intensity above the after-discharge threshold. We defined the primary visual cortex where stimulation consistently elicited a percept of a flash of light (i.e., phosphene). **Fig. 6A** presents the probability of stimulation-induced phosphene averaged across all 30 patients.

**Fig. 6.**
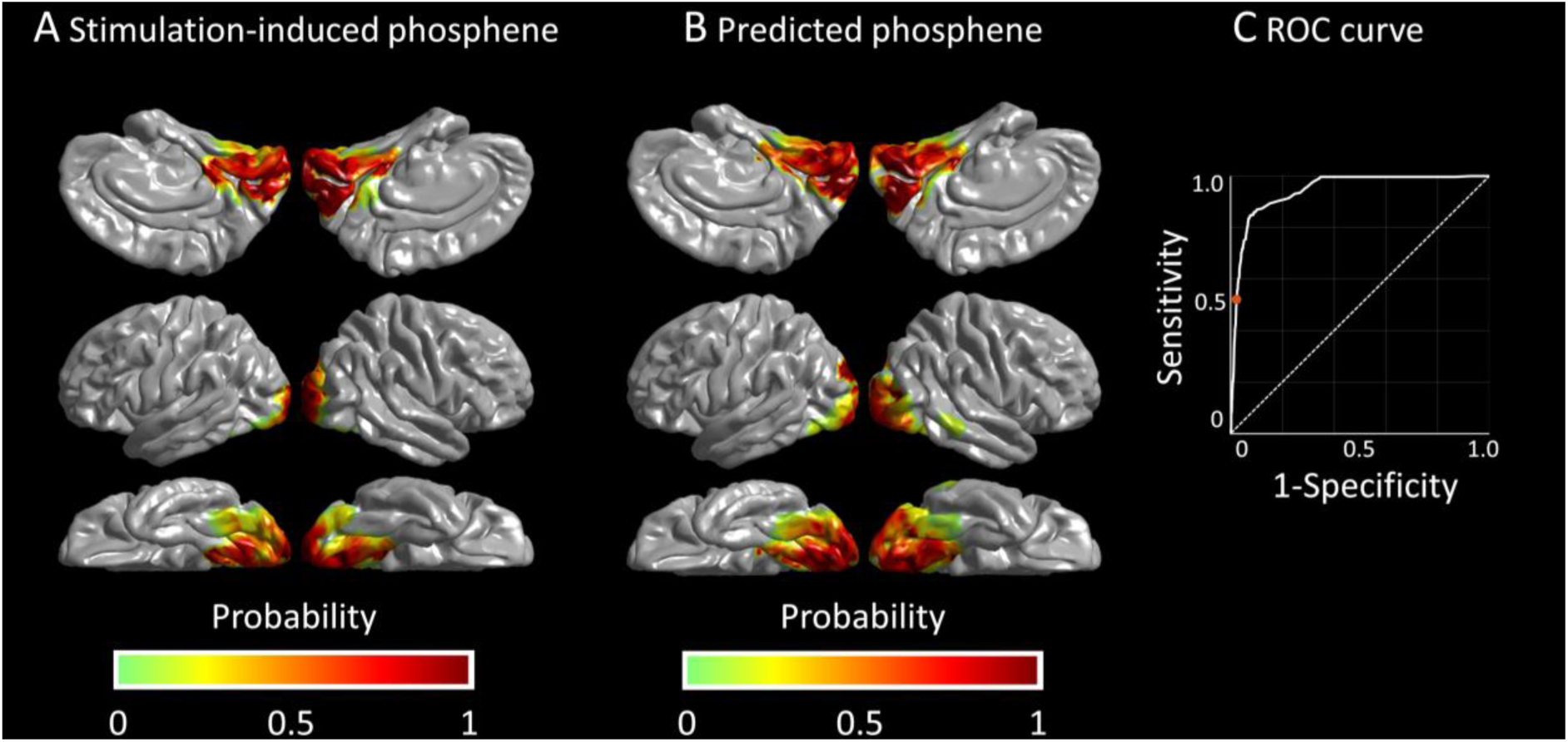
Prediction of the stimulation-defined primary visual cortex. (**A**) The spatial distribution of primary visual sites as suggested by the gold-standard electrical stimulation mapping. (**B**) The spatial distribution of primary visual sites as predicted by the bagged-tree-ensemble model. (**C**) The receiver-operating characteristics (ROC) analysis found the accuracy of bagged-tree-ensemble model to be 95%. Red filled circle: sensitivity of 52%, specificity of 98%, positive predictive value (PPV) of 57%, and negative predictive value (NPV) of 97%.

### 2.8. Statistical analysis

**[Aims 1 and 2]** The mixed model analysis tested the hypotheses that [a] increased proximity to the occipital pole and [b] saccade direction towards the sampled hemisphere would be independently associated with more intense striatal high-gamma suppression following saccade onset. The dependent variable was the striatal high-gamma amplitude change during the post-saccade 400-ms period (averaged across saccade events). This 400-ms period was unaffected by the next saccade event (see the behavioral results below). The fixed effect predictors included [a] the distance (mm) from the occipital pole, [b] ipsilateral saccade (1 if saccade direction was ipsilateral to the sampled hemisphere), [c] saccade direction (1 if saccade direction was left), [d] mean saccade duration in a given patient (ms), [e] SOZ involving the frontal lobe, [f] SOZ involving the temporal lobe, [g] SOZ involving the parietal lobe, [h] MRI lesion (1 if a lesion was present on MRI), [i] sex (1 if male), [j] patient age (years), and [k] the number of oral antiepileptic drugs (reflecting the severity of seizure burden and cognitive impairment [Kwan and Brodie, 2001]). The intercept and patient were treated as random effects. The significance level was set at *P* = 0.05 (IBM SPSS statistics 25 software; IBM Crop., Chicago, IL, USA).

**[Aim 3]** We tested the hypothesis that more intense striatal high-gamma suppression immediately *<u>preceding</u>* saccade onset would predict the upcoming saccade’s direction and duration on a trial-by-trial basis. The mixed logistic regression analysis determined whether smaller striatal high-gamma amplitude (i.e., more intense high-gamma suppression) during the 100-ms period immediately before saccade onset, as a fixed-effect predictor, would increase the probability of the upcoming saccade being directed to the ipsilateral direction. The mixed model analysis likewise determined whether smaller striatal high-gamma amplitude immediately before saccade onset, as a fixed-effect predictor, would increase the duration of the upcoming saccade. The intercept and patient were treated as random effects.

**[Aim 4]** The following two analyses determined how accurately saccade-related high-gamma modulations would localize the ESM-defined primary visual cortex. First, the mixed logistic regression analysis determined whether [a] an increase in the *<u>absolute</u>* high-gamma amplitude percent change during the post-saccade 400-ms period, [b] ipsilateral saccade (1 if saccade direction was ipsilateral to the sampled hemisphere), [c] electrodes location (1 if electrodes located within the occipital-temporal lobes), as fixed-effect predictors, would increase the probability of a given electrode site being classified as the primary visual cortex. The *<u>absolute</u>* high-gamma amplitude percent change (i.e., area under the curve) would be increased if saccades resulted in either augmentation or suppression of high-gamma activity. The intercept and patient were treated as random effects.

Subsequently, the machine learning algorithm referred to as the bagged-tree-ensemble model (Breiman, 1996, 2001) further validated the mixed model-based localization of the primary visual cortex using the aforementioned high-gamma measure. Without the investigators’ supervision, the hyperparameter optimization function, implemented in the Machine Learning and Deep Learning Toolbox (MATLAB R2020a; MathWorks Inc., Natick, MA), maximally improved the classifiers for ESM-defined primary visual sites. The hyperparameters employed in the present study included: ensemble method, maximum number of splits, number of learners, number of predictors to sample (Johnson *et al*., 2019). We used the k-fold cross-validation technique, which randomly partitioned the data into ten disjoint folds of roughly equal size (i.e., each k-th fold served as a test dataset for the model trained on the dataset excluding the k-th fold data). We evaluated the model performance of the bagged-tree-ensemble model using [a] the area under the receiver operating characteristics (ROC) curve, [b] positive and negative predictive values (PPV and NPV), and [c] the accuracy score (**Fig. 6C**). PPV (NPV) was defined as [the number of true positive (negative) sites] among [the number of predicted positive (negative) sites]. The accuracy score was defined by [the number of true positive sites plus true negative sites] divided by [the total number of electrode sites]. Based on the prediction made by the bagged-tree-ensemble model, we finally delineated the distribution of electrode sites predicted to be the primary visual cortex on the 3D brain surface (**Fig. 6B**). We have provided the **supplementary document**, including **.mat file**, which consists of our empirical bagged-tree-ensemble model allowing investigators to test and use our model for mapping the visual cortex in their own patient cohorts.

## 3. Results

### 3.1. Behavioral results

The mean number of identified saccade events in each direction was 97.9 per patient (range: 81-100 per patient). The duration of left- and right-ward saccades averaged across patients was 118.2 ms (range: 83.7-221.0 ms) and 122.5 ms (83.4-217.0 ms), respectively. The minimal interval between saccade onset and the next employed saccade event was 542 ms on average across patients (SE: 54 ms).

### 3.2. Saccade-related high-gamma patterns were similar across different analytic approaches

The Spearman’s rank test demonstrated that the spatial distribution (rated as the rank order) of high-gamma modulations was similar between the fixed and jittered baseline approaches (Spearman’s rho averaged across patients: 0.99) as well as between the fixed baseline and normalization approaches (mean Spearman’s rho: 0.94).

### 3.3. 4D visualization of peri-saccadic high-gamma modulations

Fig. 5 and **Video S1** present the 4D atlas of peri-saccadic high-gamma dynamics at the whole-brain level. The 4D atlas demonstrated a transient high-gamma suppression initially in the striatal region and subsequently in the lateral-occipital and fusiform regions following saccade onset (Fig. 5A). Following saccade offset (Fig. 5B), high-gamma suppression lingered in the lateral-occipital region, whereas high-gamma augmentation took place in the striatal and fusiform regions. Saccade-related high-gamma suppression within the striatal cortex appeared intense at sites proximal to the occipital pole.

### 3.4. Peri-saccadic high-gamma modulations in a given ROI

Fig. 4 shows the results of the ROI-based assessment of peri-saccadic high-gamma modulations. Significant high-gamma suppression took place at −90 ms to ipsilateral saccade onset in the striatal regions, +15 ms in the lateral-occipital, and −90 ms in the fusiform. Significant high-gamma suppression lasted until +80 ms after ipsilateral saccade offset in the striatal regions, +235 ms in the lateral-occipital, and +105 ms in the fusiform. Significant high-gamma augmentation took place within +125 ms after ipsilateral saccade offset in the striatal region and +235 ms in the fusiform region, whereas high-gamma augmentation did not reach significance in the lateral-occipital regions.

The FEFs contralateral to the saccade direction showed significant high-gamma augmentation by +255 ms after saccade onset, and such augmentation lasted until saccade offset (Fig. 4).

### 3.5. [Aims 1 and 2] Peri-saccadic striatal high-gamma modulations and saccade behaviors

A total of 127 striatal electrode sites were available for the mixed model analysis (mean distance from the occipital pole: 31.6 mm [SD: 12.4 mm]). The mixed model revealed that striatal high-gamma suppression during the post-saccade 400 ms period was intense at sites proximal to the pole (mixed model estimate: +0.26; 95% CI: 0.16, 0.36; t = 5.08; *P* < 0.001), ipsilateral to the direction of given saccades (mixed model estimate: −4.32; 95% CI: −6.43, - 2.20; *t* = 4.02; *P* < 0.001), and in patients whose mean saccade duration was prolonged (mixed model estimate: −0.10; 95% CI: −0.20, −0.01; *t* = −2.19; *P* = 0.036; **Table 2**).

**Table 2:** Factors associated with saccade-related high-gamma modulations.

| Variables | Estimate | SE | df | *t* | *P* | 95% CI |  |
| --- | --- | --- | --- | --- | --- | --- | --- |
|  |  |  |  |  |  | LL | UL |
| (Intercept) | 7.98 | 10.72 | 24.05 | 0.74 | 0.46 | -14.15 | 30.11 |
| Distance from the occipital pole * | 0.26 | 0.05 | 252.94 | 5.08 | <0.001 | 0.16 | 0.36 |
| Ipsilateral saccade ** | -4.32 | 1.07 | 243.06 | -4.02 | <0.001 | -6.43 | -2.20 |
| Saccade direction (left) | -0.64 | 1.08 | 245.91 | -0.60 | 0.55 | -2.77 | 1.48 |
| Saccade duration *** | -0.10 | 0.05 | 31.70 | -2.19 | 0.036 | -0.20 | -0.01 |
| Seizure onset zone |  |  |  |  |  |  |  |
| Frontal | 1.04 | 3.25 | 19.28 | 0.31 | 0.75 | -5.76 | 7.83 |
| Temporal | 4.01 | 3.56 | 18.37 | 1.13 | 0.27 | -3.45 | 11.5 |
| Parietal | -1.99 | 3.66 | 19.30 | -0.54 | 0.59 | -9.64 | 5.67 |
| MRI lesion | -4.51 | 3.56 | 18.90 | -1.26 | 0.22 | -11.99 | 2.96 |
| Sex (male) | 2.19 | 2.90 | 18.94 | 0.75 | 0.46 | -3.89 | 8.26 |
| Age | -0.28 | 0.46 | 19.77 | -0.61 | 0.55 | -1.25 | 0.68 |
| Number of AEDs | 0.74 | 2.33 | 20.40 | 0.32 | 0.75 | -4.12 | 5.61 |

AEDs = Antiepileptic drugs. CI = Confidence interval. LL = Lower limit. UL = Upper limit.
SE = Standard error. \*: Each 1-mm added distance from the occipital pole increased the amplitude of saccade-related striatal high-gamma activity by 0.26%; in other words, striatal sites more proximal to the occipital pole showed more intense saccade-related high-gamma suppression. \*\*: Saccades ipsilateral to the iEEG sampling hemisphere decreased the amplitude of saccade-related striatal high-gamma activity by 4.32%. \*\*\*: Patients with a 1-ms longer mean saccade duration had 0.1% smaller saccade-related striatal high-gamma activity.

### 3.6. [Aim 3] Preceding striatal high-gamma modulations predicted saccade behaviors

The mixed logistic regression analysis revealed that smaller preceding striatal high-gamma amplitude change (i.e., more intense high-gamma suppression) was associated with a higher probability of the upcoming saccade directed to the side ipsilateral to the sampled hemisphere (mixed model estimate: −1.02 × 10^-3^; 95% CI: −1.49 × 10^-3^, −5.42 × 10^-4^; *t* = −4.20; *P* < 0.001).

The mixed model analysis, employed to ipsilateral saccade events, revealed that smaller preceding striatal high-gamma amplitude change was associated with longer duration of the upcoming saccade (mixed model estimate: −2.07 × 10^-2^; 95% CI: −3.93 × 10^-2^, −2.04 × 10^-3^; *t* = −2.18; *P* = 0.03). Conversely, the analysis employed to contralateral saccade events failed to show a significant association between the preceding striatal high-gamma amplitude and the duration of the upcoming saccade (mixed model estimate: −1.24 × 10^-2^; 95% CI: −2.79 × 10^-2^, 3.09 × 10^-3^; *t* = −1.57; *P* = 0.12).

### 3.7. [Aim 4] Task-free high-gamma modulations localized the primary visual cortex

The mixed logistic regression analysis revealed that increased absolute saccade-related high-gamma modulation localized the primary visual cortex (mixed model estimate: +0.36; 95% CI: 0.32, 0.40; *t* = 17.57; *P* < 0.001). In other words, each 1% increase in │saccade-related high-gamma modulation│ increased the odds of a given site being the primary visual cortex by 1.43 (i.e., *e*^0.36^). The aforementioned high-gamma effect was significant independently of the anatomical location of electrode sites. Namely, compared to electrode sites within the frontal-parietal region, those within the occipital-temporal lobes (mixed model estimate: +3.80; 95% CI: 2.93, 4.68; *t* = 8.52; *P* < 0.001) had a higher odds for being classified as the primary visual cortex. The direction of saccades did not show a significant effect (mixed model estimate: +0.20; 95% CI: −0.14, 0.54; *t* = 1.13; *P* = 0.25; **Table S2**).

The mixed logistic regression model localized the primary visual cortex with an area-under-the-curve score of 0.98 (**Fig. S2**). The bagged-tree-ensemble model localized the primary visual cortex with an area-under-the-curve score of 0.94 and an accuracy of 95% (**Figs. 6B-C**).

## 4. Discussion

### 4.1. Mechanistic significance of high-gamma modulations before, during, and after saccades

The present study provided the first-ever 4D atlas animating the dynamics of peri-saccadic high-gamma modulations with a temporal resolution of 5 ms at the whole-brain level (**Video S1;** Fig. 5). Our iEEG signal sampling involved the medial and inferior occipital-temporal surfaces, which would be challenging to assess with noninvasive neurophysiology recordings alone (Fig. 1A**; Table S1**). The 4D dynamic atlas expanded our understanding of the cortical mechanism of saccadic suppression based on the previous observations of single-neuron recordings in non-human primates (Duffy and Burchfiel, 1975; Zanos *et al*. 2016; Berman *et al*., 2017) as well as iEEG recordings in small numbers of patients with focal epilepsy (Uematsu *et al*., 2013; Golan *et al*., 2017). Our atlas successfully revealed that saccade-related neural suppression involved *<u>contiguous</u>* posterior cortical networks with a spatial gradience and that such suppression was initated in the striatal cortex and rapidly involved the lateral-occipital and fusiform regions (Fig. 4). Investigation of 2,290 subdural iEEG electrodes allowed us to delineate the spatial contiguity of event-related neural modulation patterns.

**Aims 1 and 2** allowed us to better understand the saccadic suppression, a principal mechanism minimizing the perception of blurred vision during rapid eye movements. Specifically, saccade-related neuronal suppression was found intense in the posterior portion within the striatal cortex, when saccades were directed ipsilaterally to the iEEG sampling hemisphere, and in patients with a long mean duration of saccades (**Table 2**). Our study’s novelty lies in the rigorous statistical analysis. The anatomical and behavioral effects on saccade-related high-gamma measures were significant, independently of the impacts of patient demographics and epilepsy-related variables (**Table 1**). Collective evidence indicates that saccadic suppression is, at least in part, supported by transient suppression of the large-scale, posterior cortical networks, especially the primary visual cortex processing the near-foveal but less-attended hemifields. A previous study of healthy individuals using transcranial magnetic stimulation (TMS) demonstrated the causal role of the primary visual cortex in saccadic suppression (Thilo *et al*., 2004). The TMS study demonstrated that saccades did not reduce the intensity of phosphene percepts induced by magnetic excitation of the primary visual cortex.

**Aim 3** provided novel evidence that the primary visual cortex *per se* actively prepares it in advance for massive image motion expected during long saccades. Expressly, the trial-by-trial analysis, on an individual patient level, indicated that attenuated striatal high-gamma activity predicted the upcoming saccade to be directed ipsilaterally to the iEEG sampling hemisphere and to have a longer duration. Since signal sampling from the midbrain is not ethically feasible, we cannot completely rule out the possibility that the superior colliculus of the midbrain would drive the saccadic suppression at the cortical level. A single-neuron study of non-human primates previously reported that neural suppression of the superior colliculus might take place 70 ms before saccade onset (Hafed and Krauzlis, 2010). The present study revealed that significant striatal high-gamma suppression took place as early as 90 ms before the onset of ipsilateral saccades (Fig. 4A). The current iEEG study cannot directly assess the role of the thalamus in saccadic suppression. A previous study of healthy humans using functional MRI reported that saccade-related hemodynamic suppression involved both striatal cortex and thalamus; thus, the earliest site driving saccadic suppression was estimated to lie at or before the primary visual cortex (Sylvester *et al*., 2005). Continuous iEEG monitoring via deep brain stimulation therapy may provide unique opportunities to measure saccade-related thalamic modulations in the future.

The present study was characterized by the assessment of neural modulations related to saccades *<u>spontaneously</u>* occurring during *<u>task-free, naturalistic</u>* wakefulness. We found that saccade-related high-gamma augmentation was minimal in the lateral-occipital region. This observation contrasts with our previous iEEG report that intense high-gamma augmentation involved the lateral-occipital regions during picture naming task, in which participants needed to attend a series of visual stimuli (Nakai *et al*., 2019). In the present study, significant striatal high-gamma augmentation took place bilaterally within 125 ms after the saccade offset (Fig. 4A); this neural activation is primarily attributed to the retinal image updated at a given saccade offset. We did not find evidence that the FEF initiated saccadic suppression. Significant FEF high-gamma augmentation was noted only in trials when saccades were directed contralaterally to the iEEG sampling hemisphere, and such neuronal activation took place >200 ms after saccade onset (Fig. 4D).

### 4.2. Clinical significance of saccade-related high-gamma modulations during task-free wakefulness

**Aim 4** provided evidence that high-gamma modulations associated with spontaneous saccades help localize the primary visual cortex. The mixed logistic regression analysis revealed that each 1% increase in the *<u>absolute</u>* saccade-related high-gamma modulation increased the odds of a given site being the ESM-defined primary visual cortex by 1.43. In other words, both augmentation and attenuation of high-gamma amplitude provided information useful to localize the visual cortex. The machine learning-based analysis likewise localized the primary visual cortex with an area-under-the-curve score of 0.94 and an accuracy of 95% (**Figs. 6B-C**). Measurement of spontaneous event-related high-gamma modulations may be useful in the evaluation of patients uncooperative to ESM. Readers are welcome to externally validate our model incorporated in the **Supplementary .mat file** (see the online supplementary document).

Our study has provided an analytic approach to identify the broadband high-frequency activity of physiological nature. Investigators have hypothesized that spontaneous broadband high-frequency activity at >40 Hz may serve as a biomarker of the epileptogenic zone (Cimbalnik *et al*., 2018; Jobst *et al*., 2020; Avigdor *et al*., 2021). Yet, the distinction between physiological and pathological ones could be challenging without behavioral monitoring (Nagasawa *et al*., 2012; Matsumoto *et al*., 2013; Nonoda *et al*., 2016). The current study suggests that the assessment of the iEEG-EOG temporal relationship may reduce the risk of misinterpreting some of the physiological high-frequency activities as pathologic ones. A previous iEEG study reported that external sounds augmented high-gamma activity in the auditory cortex even during task-free wakefulness (Arya *et al*., 2018*b*). Further studies are warranted to determine what modality should be monitored to optimize the diagnostic utility of spontaneous high-frequency activity for localization of the epileptogenic zone.

### 4.3. Methodological considerations

Since the eye tracking was not employed in the present study, it remains uncertain what visual scene/object each patient fixated at a given saccade offset. Moreover, the intensity of room light was not strictly controlled across the study patients. Previous iEEG studies reported that different physical properties of visual objects accounted for different degrees of event-related high-gamma augmentation in the lower- and higher-order visual pathways (Matsuzaki et al., 2015). Even this limitation taken into account, we cannot discount the clinical utility of our iEEG-based model because it localized the primary visual cortex with an accuracy of 95% (Fig. 6C). It is infeasible to attribute the observation of striatal high gamma suppression preceding saccade onset to our event marking approach (i.e., visual marking, which may not have a millisecond precision). We found that striatal high gamma suppression occurred 90 ms before ipsilateral saccades onset (Fig. 4A).

As shown in **Videos S2 and S3**, saccade-related alpha/beta suppression took place together with high-gamma augmentation, whereas alpha/beta augmentation coupled with high-gamma suppression. This qualitative observation was concordant with those reported in many other iEEG studies of event-related spectral responses (Miller *et al*., 2007; Crone *et al*., 2011; Uematsu *et al*., 2013; Nakai *et al*., 2017). Because a single high-gamma cycle is much shorter in duration than alpha/beta ones, the rapid neural dynamics may be better delineated with high-gamma measures.

The present study did not reveal a significant age effect on the peri-saccadic high-gamma modulations. Since we did not include patients younger than five years of age, the developmental changes of peri-saccadic high-gamma modulations during infancy and toddlerhood remain to be determined by future studies.

## Abbreviations

EOG: electrooculography
ESM: electrical stimulation mapping
4D: four-dimensional
FEF: frontal eye field
FIR: finite impulse response
HFO: high-frequency oscillation
iEEG: intracranial EEG
NPV: negative predictive value
PPV: positive predictive value
ROC: receiver operating characteristics
ROI: region of interest
SOZ: seizure onset zone
3D: three-dimensional
TMS: transcranial magnetic stimulation.

## Acknowledgments

We are grateful to Karin Halsey, BS, REEGT. and Jamie MacDougall, RN, BSN, CPN at Children’s Hospital of Michigan for the collaboration and assistance in performing the studies described above. This work was supported by NIH grant NS064033 (to E.A.).

## Figure Legends

We have embedded all figures and figure legends within the main text. The same figures are also provided as separate TIFF files.

