## supplemental documents for "Spontaneous modulations of high frequency cortical activity"

Eishi Asano [*]

$: Equal contribution.

**Figure S1: Saccade-related high-gamma modulations quantified using the three different approaches.**

**Figure S2: Utility of saccade-related high-gamma modulations in prediction of the primary visual areas.**

**Table S1: The number of electrodes at region of interests (ROIs).**

**Table S2: Mixed logistic regression model-based prediction of the primary visual cortex.**

**Video S1: Four-dimensional atlas of peri-saccadic high-gamma modulations.**

**Video S2: Four-dimensional atlas of peri-saccadic alpha modulations.**

**Video S3: Four-dimensional atlas of peri-saccadic beta modulations.**

**Supplementary .mat file: The bagged-tree-ensemble model for prediction of the primary visual areas.**

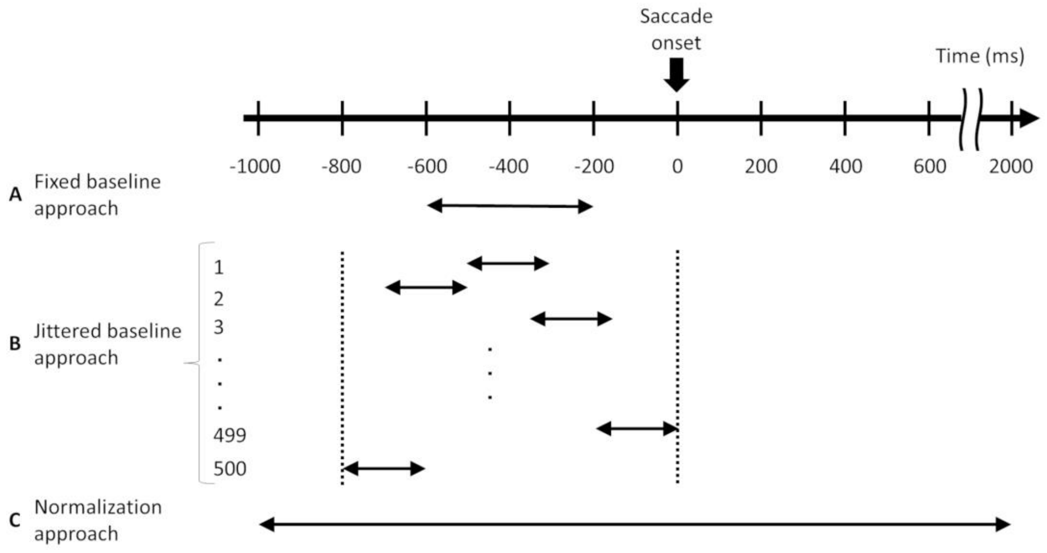


**Fig. S1. Saccade-related high-gamma modulations quantified using the three different approaches.** (**A**) [Fixed baseline approach] The baseline was defined as the average across high-gamma amplitudes during the 400-ms period 200-600 ms before saccade onset. (**B**) [Jittered baseline approach] The baseline was defined as the average across high-gamma amplitudes during a randomly-selected 200-ms period between saccade onset and 800 ms before saccade onset. (**C**) [Normalization approach] The baseline period was defined as the entire 3000-ms period between 1000 ms before and 2000 ms after saccade onset. At a given 5-ms time window and each channel, we computed 'high-gamma z-score', reflecting how much high-gamma amplitude deviated from the mean during the baseline period.

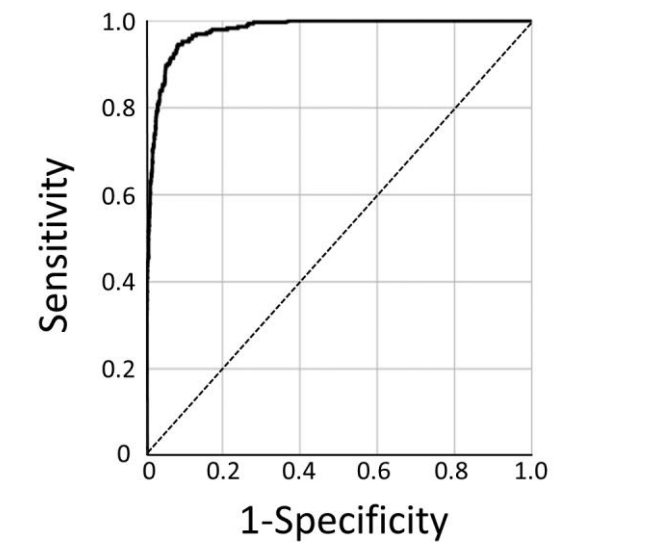


**Fig. S2. Mixed logistic regression model-based prediction of the primary visual cortex.**

The receiver-operating characteristics (ROC) analysis indicated the accuracy of the mixed logistic regression model for localization of the primary visual cortex to be an area under the curve (AUC) of 0.98.

**Table S1: The number of electrodes at region of interests (ROIs).**

| ROIs | Left | Right | Total |
|---|---|---|---|
| superior frontal gyrus (SFG) | 42 | 58 | 100 |
| rostral middle-frontal gyrus (rMFG) | 58 | 97 | 155 |
| caudal middle frontal gyrus (cMFG) | 61 | 75 | 136 |
| pars orbitalis of the inferior-frontal gyrus (POr) | 14 | 20 | 34 |
| pars triangularis of the inferior-frontal gyrus (PTr) | 39 | 60 | 99 |
| pars opercularis of the inferior-frontal gyrus (Pop) | 51 | 23 | 74 |
| frontal-eye field (FEF) | 40 | 37 | 77 |
| medial orbitofrontal gyrus (MOrb) | 3 | 1 | 4 |
| lateral orbitofrontal gyrus (LOrb) | 5 | 21 | 26 |
| paracentral gyrus (PCL) | 8 | 13 | 21 |
| precentral gyrus (PreCG) | 83 | 121 | 204 |
| postcentral gyrus (PoCG) | 111 | 103 | 214 |
| precuneus (PCun) | 11 | 20 | 31 |
| superior parietal lobule (SPL) | 17 | 12 | 29 |
| supramarginal gyrus (SMG) | 80 | 87 | 167 |
| inferior parietal lobule (IPL) | 21 | 34 | 55 |
| superior-temporal gyrus (STG) | 87 | 108 | 195 |
| middle-temporal gyrus (MTG) | 77 | 53 | 130 |
| inferior-temporal gyrus (ITG) | 43 | 45 | 88 |
| fusiform gyrus (FG) | 48 | 56 | 104 |
| temporal pole (TP) | 0 | 1 | 1 |
| entorhinal gyrus (Ent) | 3 | 6 | 9 |
| parahippocampal gyrus (PHG) | 7 | 9 | 16 |
| lateral-occipital gyrus (LOG) | 69 | 71 | 140 |
| anterior cingulate gyrus (aCG) | 1 | 5 | 6 |
| posterior cingulate gyrus (pCG) | 19 | 29 | 48 |
| striatal | 60 | 67 | 127 |

**Table S2: Mixed logistic regression model-based prediction of the primary visual cortex.**

| | | | | | | 95% CI | |
|---|---|---|---|---|---|---|---|
| Variables | Estimate | SE | df | *t* | *P* | LL | UL |
| (Intercept) | -9.16 | 0.61 | 4576 | -14.96 | <0.001 | -10.36 | -7.96 |
| Saccade-related │high-gamma (%)│ | 0.36 | 0.02 | 4576 | 17.57 | <0.001* | 0.32 | 0.40 |
| Ipsilateral saccade | 0.20 | 0.17 | 4576 | 1.13 | 0.25 | -0.14 | 0.54 |
| Electrode location | 3.80 | 0.45 | 4576 | 8.52 | <0.001 | 2.93 | 4.68 |

CI = Confidence interval. LL = Lower limit. UL = Upper limit. *: Each 1% increase in the absolute saccade-related high-gamma modulation increased the odds of a given site being the primary visual cortex by 1.43 (i.e., $e^{0.36}$).

**Video S1: Four-dimensional atlas of peri-saccadic high-gamma modulations.**

The 4D atlas demonstrated a transient high-gamma suppression initially in the striatal region and subsequently in the lateral-occipital and fusiform regions following saccade onset. Subsequently, high-gamma suppression lingered in the lateral-occipital region, whereas high-gamma augmentation took place in the striatal and fusiform regions. Saccade-related high-gamma suppression within the striatal cortex appeared intense at sites proximal to the occipital pole.

**Video S2: Four-dimensional atlas of peri-saccadic alpha modulations.**

The 4D atlas demonstrated a transient alpha augmentation in the occipital regions following saccade onset. Subsequently, alpha attenuation took place in the striatal regions, whereas alpha augmentation lingered in the occipital pole, lateral-occipital, and fusiform region.

**Video S3: Four-dimensional atlas of peri-saccadic beta modulations.**

The 4D atlas demonstrated a transient beta augmentation in the occipital regions following saccade onset. Subsequently, beta attenuation took place in the striatal regions, whereas beta augmentation lingered in the occipital pole, lateral-occipital, and fusiform region.

**Supplementary .mat file: The bagged-tree-ensemble model for prediction of the primary visual sites using saccade-related high-gamma activity.**

This MATLAB-based .mat file contains [1] 'trainedModel' including our bag-tree-ensemble model and [2] 'Dataset' from 2,290 nonepileptic electrode sites of all 30 study patients. 'Explanatory_var': │high-gamma amplitude (%)│ during ipsilateral and contralateral saccades recorded at 2,290 nonepileptic sites (i.e., 4,580 measures). 'Elec_T_O' of 1: Electrode sites within the occipital-temporal lobe. 'Saccade_electrode_correlation' of 1: Ipsilateral saccade. 'Response_var' of 1: Primary visual site defined by electrical stimulation mapping.

To validate our bag-tree-ensemble-based prediction, users will run the following statement within the MATLAB command window:

```
yfit = trainedModel.predictFcn(Dataset)
```

Users will find the list of model-predicted primary visual sites ('yfit' of 1: primary visual sites based on the prediction model).

To predict the primary visual sites on a new iEEG dataset, users will run the following statement within the MATLAB command window:

```
yfit = trainedModel.predictFcn(T)
```

Thereby, 'T' (table format) will need to have the three variables, including 'Explanatory_var' (double format), 'Elec_T_O' (categorical), and 'Saccade_electrode_correlation' (categorical).
